# Coconut filters woven using Native Hawaiian techniques capture coastal microplastic and sunscreen pollution and support fungal bioremediation

**DOI:** 10.64898/2026.09.05.749617

**Authors:** Vera H. Wang, Kaylee E. Christensen, Anthony Amend

## Abstract

Microplastic pollution and sunscreen contaminants increasingly threaten coastal ecosystems, yet many existing clean-up solutions rely on synthetic materials, creating additional carbon emissions during manufacturing, or address pollutant removal without subsequent treatment, leaving risk of future run-off back into the ocean. Here we evaluate coconut husk, an abundant and biodegradable agricultural byproduct, as a dual-function platform for physical pollutant capture and microbial bioremediation applications. To test the natural microbial community of coconuts for bioremediation potential, we isolated fungi from coconut husks and screened their growth under plastic- and sunscreen-containing conditions. 24 fungal isolates grew on plastic- and/or sunscreen media, with several isolates belonging to genera previously documented to degrade plastic, i.e. *Chaetomium*, *Curvularia*, *Meyerozyma*, *Cladosporium*, and *Fusarium*. We then developed filtration modules using coconut husks, fiber, and leaves, basing our design on traditional Hawaiian pāpale lau niu (coconut leaf hat) weaving. In filtration trials, the woven devices captured up to 76.58 ± 3.73% of suspended microplastic particles in 2 minutes and captured suspended sunscreen with a 533.51 ± 0.81 NTU decrease in net turbidity of the surrounding water than in control trials with no filter. Overall, this work introduces a form of natural innovation for pollution clean-up, building on coconut-based materials, traditional techniques, and microbial resilience to create a sustainable and cost-effective system.

## Introduction

Each year an estimated 14,000 tons of sunscreen wash into the ocean (Shaath, 2005) while more than 170 trillion plastic particles, totaling 1.1–4.9 million tonnes, persist at the sea surface (Eriksen et al., 2023). Both pollutants pose their own ecological threats. Sunscreen-derived chemicals disrupt nutrient cycling (Rodríguez-Romero et al., 2019) and exhibit toxicity across marine organisms (Lozano et al., 2020). Plastic particles likewise endanger marine macrobiota and interfere with cellular processes (Gregory, 2009; Bucci, Tulio & Rochman, 2020; Mason et al., 2022). Together these pollutants create additional compounded threats as sunscreen accumulates on lipophilic plastic particles, causing increased dispersal of sunscreen and decreased activity of beneficial microbes on the plastic surface (Lee et al., 2025). Coastal environments such as coral reefs which receive constant direct anthropogenic interaction are particularly at risk (Sánchez-Quiles, Blasco & Tovar-Sánchez, 2020; Thushari & Senevirathna, 2020; Downs, Cruz & Remengesau Jr., 2022; Pinheiro et al., 2023), where microplastics can impair coral photosynthetic performance, metabolism, and calcification (Lanctôt et al., 2020; Mendrik et al., 2021), while sunscreen-derived UV filters may act as additional local stressors under high exposure and elevated-temperature conditions (Downs et al., 2002; Breakell et al., 2024). The combined ecosystem-level damage and downstream human health risks invoke immediate need for widespread remediation solutions.

Current pollutant clean-up approaches tend to focus solely on plastic removal, and many have been inefficient or caused unwanted negative environmental impacts. Mechanical ocean-debris removal systems often trap marine life as bycatch or even release more waste through machine breakdown (Falk-Andersson et al., 2023). Furthermore, many of these mechanical solutions are constructed from synthetic materials such as styrofoam, polyvinyl chloride, and high-density polyethylene (HDPE) (Nikiema & Asiedu, 2022), adding the environmental costs associated with plastic production and end-of-life disposal. These artificial devices can ultimately deteriorate into microplastics themselves or further contribute to landfill waste, thereby exacerbating the very issue they aim to mitigate. This is particularly relevant as an estimated 80% of marine plastic pollution originates from land-based sources, resulting from mismanaged waste and leakage from waste-disposal systems (Wang et al., 2024). While at least several attempts have been made for plastic removal, sunscreen remediation efforts receive far less attention. Current mitigation strategies largely emphasize the use of environmentally safer sunscreen formulations and improved wastewater treatment, as conventional treatment processes often incompletely remove organic UV filter compounds (Damikouka, Anastasopoulou & Vgenopoulou, 2024). While these efforts prevent future sunscreen pollution, they don’t address the sunscreen already in our waters. An effective, environmentally friendly, and minimally invasive clean-up strategy targeting both plastic and sunscreen pollution is thus much needed.

Pollutant removal is only the first step of clean-up. Once captured through any filtration process, pollutants are typically dumped back into landfills, perpetuating the problem of waste build up. This limitation suggests that filtration alone is an incomplete solution unless captured pollutants can be immobilized, degraded, or transformed into less harmful products. Bioremediation is the process by which microorganisms are used to fulfill this need, where the innate complex metabolisms of certain species are utilized to break down pollutants. Some microorganisms have been previously shown to transform or mineralize oxybenzone and octinoxate, the primary toxic UV filter compounds in sunscreen (Nurtayeva et al., 2025), and others can initiate the oxidation and depolymerization of synthetic plastics, turning them into more bioavailable compounds (Johnson, 2024). Fungi are particularly promising in bioremediation as many species secrete extracellular enzymes capable of attacking chemically resistant substrates like plastic (Ekanayaka et al., 2022; Steinbach, Whitner & Amend, 2025; Berger et al., 2026). An integrative approach utilizing the natural degradation abilities of microbes with environmentally-friendly filter devices could provide a practical solution to pollution remediation.

Coconut husk, the fibrous outer mesocarp of *Cocos nucifera* (coconuts), and its separated coir fibers, are lignocellulosic byproducts of coconuts that combine biodegradability, renewability, and sustainability as a promising material for environmental remediation. Coconut fiber has been tested as a filtration and absorption medium for microplastic removal from wastewater, with biofilm-enhanced fiber showing higher adsorption efficiency than untreated coir across different plastic types and particle sizes (Zharkenov et al., 2024). Coconut fiber has also been used to remove inorganic contaminants from landfill leachate, supporting its potential as a residual biomass material for treating complex contaminated effluents (Lima et al., 2025). Beyond their use in microplastic and metal removal, coconut-based materials have been incorporated into wastewater-treatment systems as low-cost adsorbent media for dye removal, owing to their favorable surface chemistry and porous structure (Chong & Tam, 2020). More recently, raw coconut fibers have been evaluated for marine oil spill remediation, where their lignocellulosic structure, high lignin content, buoyancy, and hydrophobic interactions supported crude oil adsorption under laboratory, mesoscale, and field conditions (Cardoso et al., 2025). In addition to its demonstrated effectiveness in remediation applications, coconut fiber remains an underutilized agricultural byproduct in many coconut-producing regions. After the shell, meat, and coconut water are removed, the outer husk is often burned or discarded, resulting in around 20 million tons of environmental pollution and resource waste annually (Pogosa, 2018; Stelte et al., 2022). Leveraging waste coconut husks for environmental remediation could thus provide a cost-effective and sustainable clean-up strategy.

Here we propose a design of biodegradable filter contraptions made out of coconut husks, coconut fiber, coconut leaves, and jute twine. In addition, we test whether resident fungal communities associated with coconut husks exhibit differing capacities to grow under sunscreen- and plastic-containing conditions. Our filter design aims to be easily deployable, biodegradable, and cost-effective, while also providing a substrate for plastic- and sunscreen-degrading fungi to eliminate captured particles. The design of the filtration modules are modified from the intricate patterns of the Hawaiian pāpale lau niu—translated as “coconut leaf hat” in □Ōlelo Hawai‘i. In using the pāpale lau niu design we aim to honor the Native Hawaiian traditions of sustainability while utilizing the widely known weaving technique for easy deployment. With traces back to the first Polynesian settlers of Hawai‘i, the making and use of woven items began to decline toward the end of the nineteenth century as Hawai‘i experienced increasing influence from the growing non-Hawaiian population. However, weaving lau (leaves) into pāpale (hats) persists to the modern day, providing sun protection, cultural connection, and income for local families (Keawe, MacDowell & Dewhurst, 2014). We build off these traditional weaving techniques in our design to create a culturally grounded, scalable, low-cost, and sustainable remediation strategy to reduce the persistence of microplastic and sunscreen-derived pollutants in coastal aquatic environments.

## Methods

### 1. Screening of plastic and sunscreen-tolerant fungi from coconut husks

#### 1.1 Isolation of fungi from coconut husks

For preparation of the coconut husk for isolation plates, 10 cm sections of coconut husk were removed from freshly halved coconuts using clean pliers and transferred to sterile 50 mL tubes each containing 20 mL of filter-sterilized water. Coconut husk samples were submerged in the tubes for 10 days to allow water-soluble compounds to leach into the solution. After incubation each tube was vortexed for 1 minute to dislodge the fungi from the coconut husk into suspension, creating what we refer to as the coconut husk extract. Extracts were categorized according to their visually distinct coloration as shown in Supplemental figure 2: light amber/brown and dark amber/brown. Two replicate extracts were prepared for each category.

#### 1.2 Growth on plastic and sunscreen media

The coconut husk extracts were then plated on the sunscreen and plastic media to isolate fungi tolerant of either substrate, as shown in Figure 1, with the following procedure. Initial basal growth media was prepared with 0.5% yeast extract (VWR®), 1% peptone (VWR®), and 1% agar (VWR®) for plates. 100 µg/mL chloramphenicol was added after autoclaving to prevent bacterial growth. Media was then supplemented with either 4% sunscreen (Coppertone) or 1% polyurethane (Impranil) to create the sunscreen and plastic plates. Sunscreen and polyurethane were added under laminar flow hoods after autoclaving the media to prevent degradation of either addition by heat. Non-inoculated plates served as experimental controls and were monitored throughout the experiment to ensure lack of contamination. 100 µL coconut husk extract from each tube was then plated in duplicate onto the sunscreen, plastic, and basal growth media and spread evenly. The plates were incubated for 20 days at ambient temperature. Subsequent colonies on sunscreen or plastic plates were transferred to YPD plates (1% yeast extract, 2% peptone, 2% glucose, and 1% agar), and grown at room temperature for 7 days before storage at 4°C.

**Figure 1.**
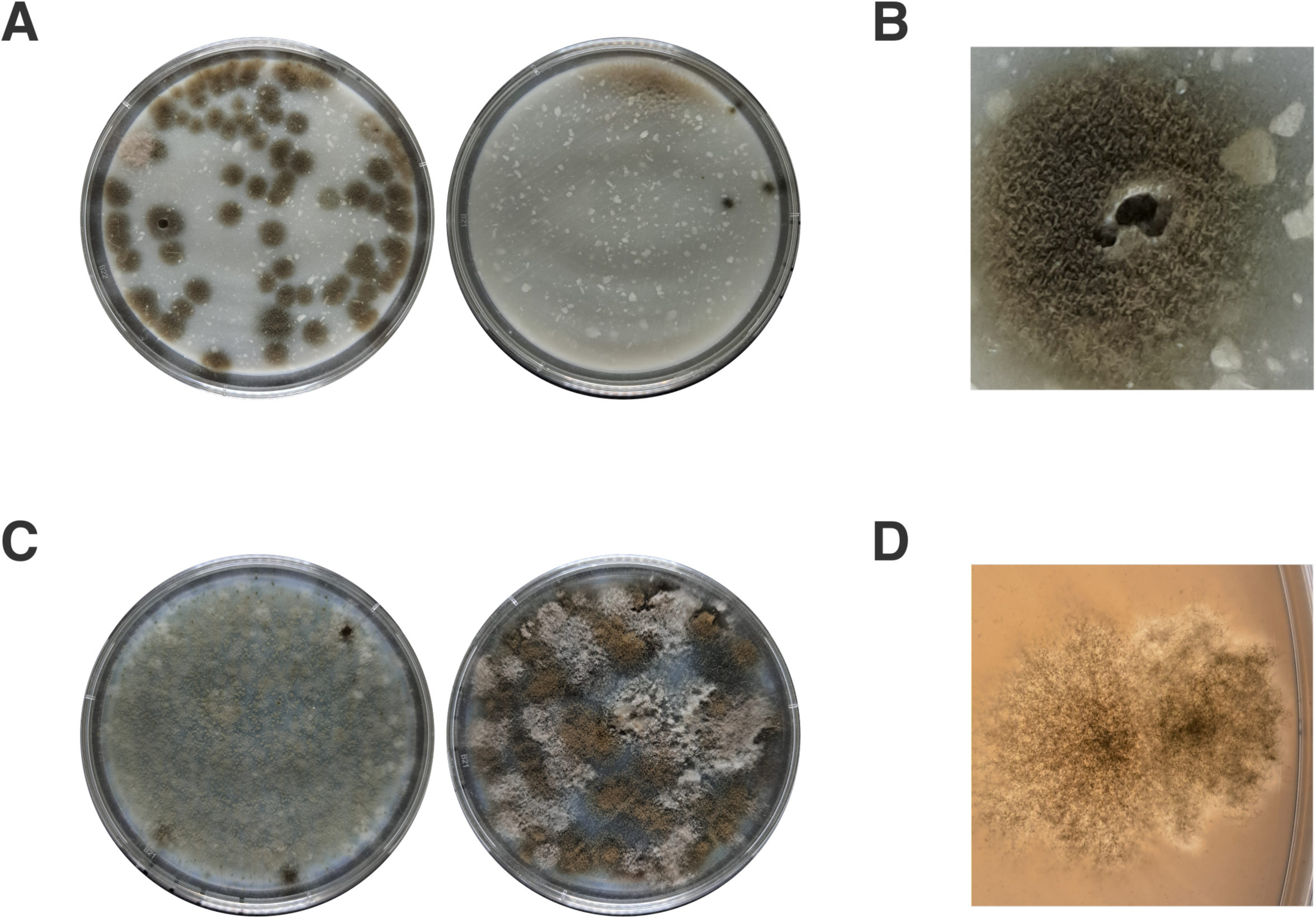
Microbial communities isolated from coconut-husk extract show robust growth patterns on plastic- and sunscreen-containing media. **A)** Coconut-husk extract grown on sunscreen media. **B)** Zoomed image of fungi interaction with sunscreen. **C)** Coconut-husk extract grown on plastic media. **D)** Zoomed image of zone of clearance around fungal growth on the plastic plate.

To test growth on sunscreen and plastic saline liquid media as in Figure 2, isolates were grown as follows. Two colonies per isolate were picked from stored YPD plates and transferred to reduced-nutrient saltwater wells containing 500 µL of 0.1% yeast extract, 0.2% peptone, 3.5% NaCl, 100 µg/mL chloramphenicol, and either 4% sunscreen (Coppertone) or 1% polyurethane (Impranil). The well plates were then grown at room temperature for 2 weeks. After incubation, 100 µL from each well was serially diluted to 10^-7^ and 5 µL of each dilution was plated onto YPD plates. After incubation for 5 days, the resulting CFUs were counted and CFUs/mL were calculated. Significant differences in mean values were analyzed using two-tailed Mann-Whitney U tests.

**Figure 2.**
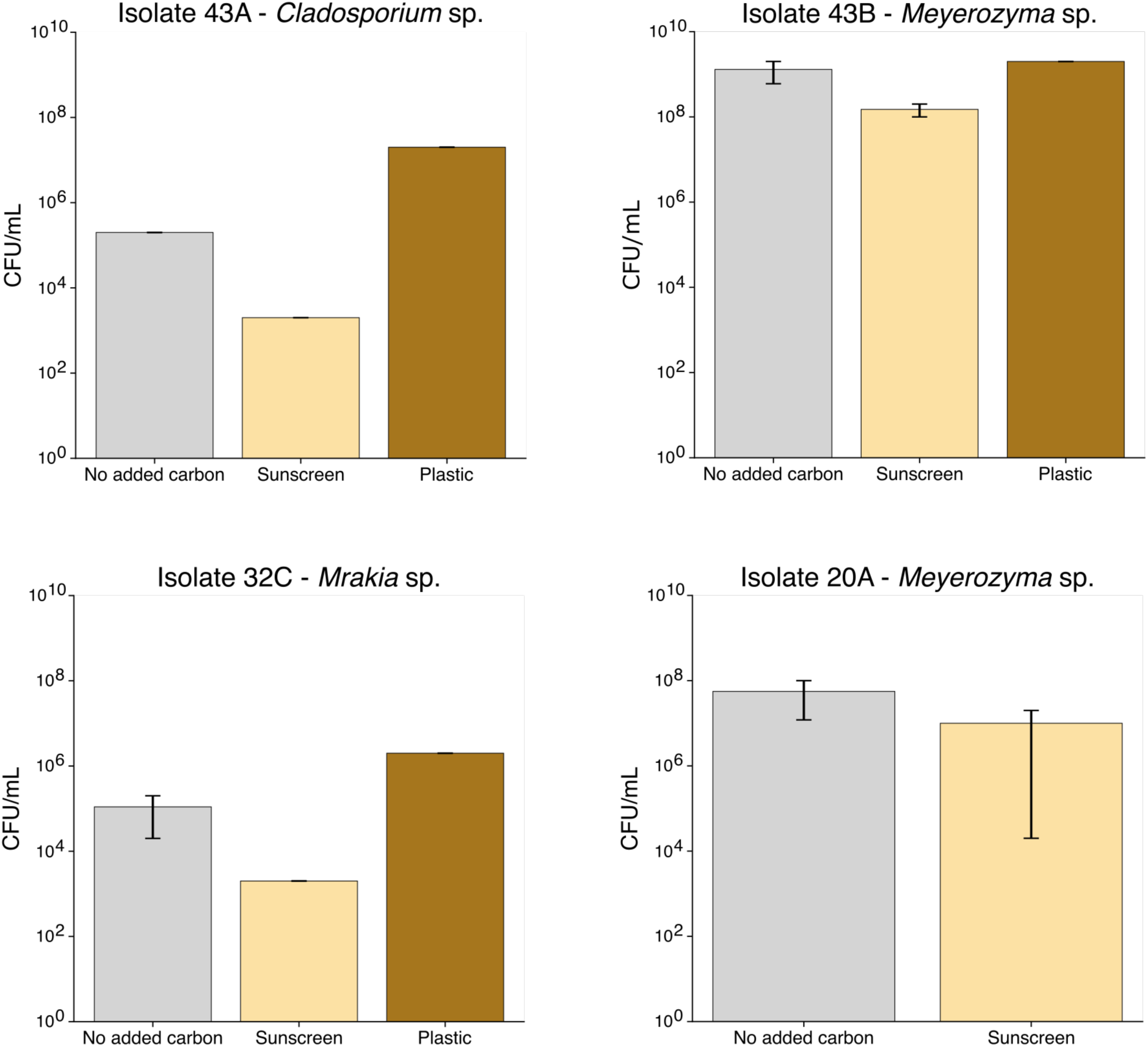
Fungal isolates sustain growth with sunscreen or plastic in carbon-limited saline liquid media. Growth of four fungal isolates in control, sunscreen-containing, and plastic-containing carbon-limited liquid media with 35 g/L NaCl, quantified as colony-forming units (CFU) per mL after seven days of growth. Isolates 43A, 43B, 20A, and 32C represent individual fungal cultures recovered from coconut husks, with their closest ITS match indicated as in Table 1. The x-axis denotes the supplement added to the carbon-limited saline media. Bars represent mean CFU/mL from n=2 replicates, with error bars indicating standard error.

**Table 1.**
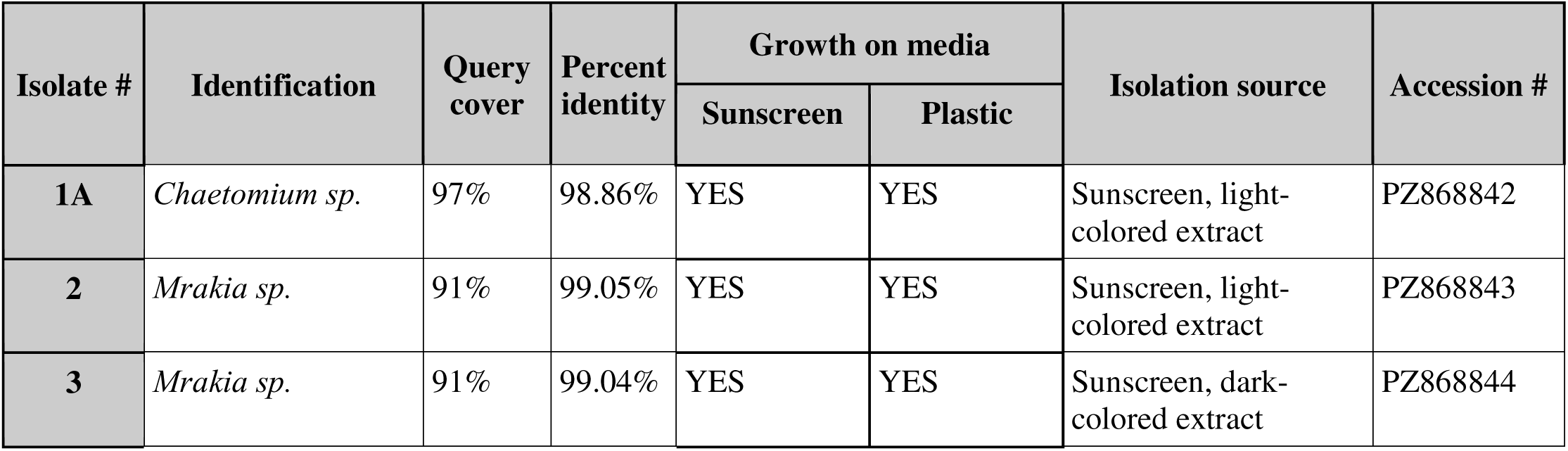
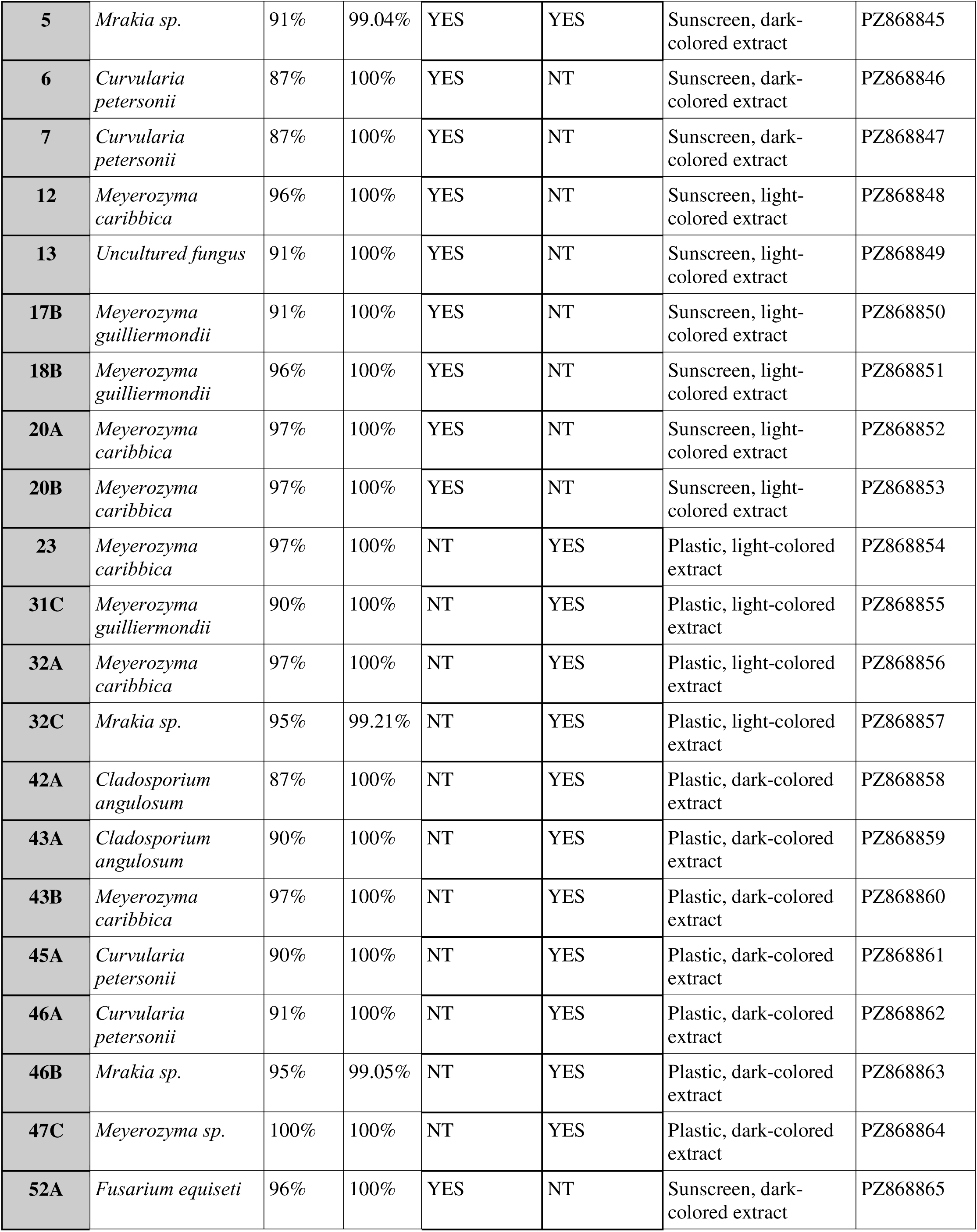
Taxonomic identification of sunscreen and plastic degrading fungi. The isolate number corresponds to the given number to which we refer to each isolate throughout the paper. The identification column indicates the species or genus of the most probable match of each isolate’s ITS sequence to the NCBI BLAST nt database. Query cover indicates the percentage of each isolate’s ITS sequence was included in the match. Percent identity indicates how similar the aligned part of each ITS sequence is to the database sequence. Growth on plastic or sunscreen media indicates that isolate’s ability to grow on either media, with NT denoting where isolates were Not Tested for the indicated treatment in this experiment. Isolation source indicates from which experimental plate the fungus was isolated from in this experiment, either on plastic or sunscreen plates and from light or dark colored coconut husk extract. Accession number refers to the associated GenBank accession for each sequence under the submission SUB16429948.

| Isolate # | Identification | Query cover | Percent identity | Growth on media |  | Isolation source | Accession # |
| --- | --- | --- | --- | --- | --- | --- | --- |
|  |  |  |  | Sunscreen | Plastic |  |  |
| 1A | *Chaetomium sp.* | 97% | 98.86% | YES | YES | Sunscreen, light-colored extract | PZ868842 |
| 2 | *Mrakia sp.* | 91% | 99.05% | YES | YES | Sunscreen, light-colored extract | PZ868843 |
| 3 | *Mrakia sp.* | 91% | 99.04% | YES | YES | Sunscreen, dark-colored extract | PZ868844 |
| **5** | *Mrakia sp.* | 91% | 99.04% | YES | YES | Sunscreen, dark-colored extract | PZ868845 |
| **6** | *Curvularia petersonii* | 87% | 100% | YES | NT | Sunscreen, dark-colored extract | PZ868846 |
| **7** | *Curvularia petersonii* | 87% | 100% | YES | NT | Sunscreen, dark-colored extract | PZ868847 |
| **12** | *Meyerozyma caribbica* | 96% | 100% | YES | NT | Sunscreen, light-colored extract | PZ868848 |
| **13** | *Uncultured fungus* | 91% | 100% | YES | NT | Sunscreen, light-colored extract | PZ868849 |
| **17B** | *Meyerozyma guilliermondii* | 91% | 100% | YES | NT | Sunscreen, light-colored extract | PZ868850 |
| **18B** | *Meyerozyma guilliermondii* | 96% | 100% | YES | NT | Sunscreen, light-colored extract | PZ868851 |
| **20A** | *Meyerozyma caribbica* | 97% | 100% | YES | NT | Sunscreen, light-colored extract | PZ868852 |
| **20B** | *Meyerozyma caribbica* | 97% | 100% | YES | NT | Sunscreen, light-colored extract | PZ868853 |
| **23** | *Meyerozyma caribbica* | 97% | 100% | NT | YES | Plastic, light-colored extract | PZ868854 |
| **31C** | *Meyerozyma guilliermondii* | 90% | 100% | NT | YES | Plastic, light-colored extract | PZ868855 |
| **32A** | *Meyerozyma caribbica* | 97% | 100% | NT | YES | Plastic, light-colored extract | PZ868856 |
| **32C** | *Mrakia sp.* | 95% | 99.21% | NT | YES | Plastic, light-colored extract | PZ868857 |
| **42A** | *Cladosporium angulosum* | 87% | 100% | NT | YES | Plastic, dark-colored extract | PZ868858 |
| **43A** | *Cladosporium angulosum* | 90% | 100% | NT | YES | Plastic, dark-colored extract | PZ868859 |
| **43B** | *Meyerozyma caribbica* | 97% | 100% | NT | YES | Plastic, dark-colored extract | PZ868860 |
| **45A** | *Curvularia petersonii* | 90% | 100% | NT | YES | Plastic, dark-colored extract | PZ868861 |
| **46A** | *Curvularia petersonii* | 91% | 100% | NT | YES | Plastic, dark-colored extract | PZ868862 |
| **46B** | *Mrakia sp.* | 95% | 99.05% | NT | YES | Plastic, dark-colored extract | PZ868863 |
| **47C** | *Meyerozyma sp.* | 100% | 100% | NT | YES | Plastic, dark-colored extract | PZ868864 |
| 52A | *Fusarium equiseti* | 96% | 100% | YES | NT | Sunscreen, dark-colored extract | PZ868865 |

#### 1.3 DNA extraction and sequencing

To identify the isolates, DNA was extracted by mixing a colony pick with 20 µL PrepMan Ultra (Applied Biosystems) and boiled at 98°C for 10 minutes. 1 µL of the supernatant was used as the template for amplification of the internal transcribed spacer (ITS) region with primers ITS1F and ITS2 (White et al., 1990). Amplicons were sequenced with Sanger sequencing and compared against the NCBI nt database using BLASTn (Altschul et al., 1990). Final sequences are available under GenBank submission SUB16429948.

### 2. Testing the woven filter devices

#### 2.1 Construction of the Baseline Prototype

The “Baseline Prototype” in Figure 3A features dried coconut fiber encased by dried coconut husks with the outer skin intact and residual coir remaining attached, constructed as follows. Two green coconuts (≈ 22 cm × 15 cm × 15 cm) were halved longitudinally with a hacksaw. After draining the coconut water, the meat and shell were scraped out and removed. Portions of the fibrous husk material were manually separated from the outer husk sections and shredded into finer strands to produce the loose coconut fiber used as the filtration medium. A 38 mm opening was drilled at each pole of every remaining husk section, and residual pith was trimmed. Both the husk sections and separated coconut fibers were sun-dried for 14 days. Once dry, three vertical half-husks were stitched into a sphere, using 3 mm holes spaced 2 cm apart around each rim. Two meters of jute twine were woven through opposing holes and tied off, producing a porous, buoyant capsule. The coconut husk capsule was then filled with 5 g of the dried coconut fiber. Photographic steps of the module construction are provided in Supplementary Data 1.

**Figure 3.**
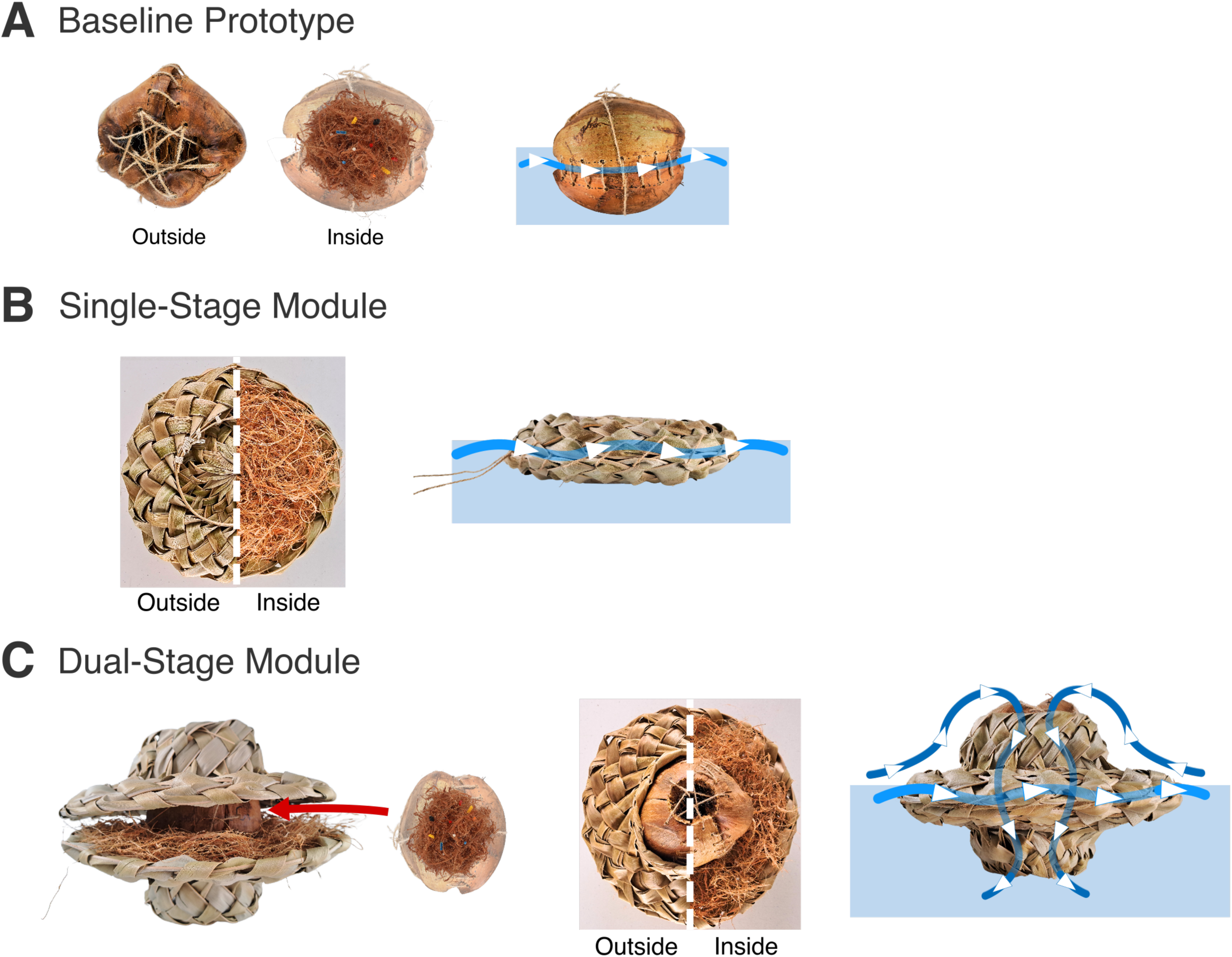
Diagrams and use of the three prototype coconut-fiber filter contraptions. **A)** Left, exterior and interior side views of the Baseline Prototype. Right, example of water flow during deployment. **B)** Left, top view and cross section of the Single-Stage Module filter featuring coconut fiber encased by pāpale lau niu brims. Right, example of water flow during deployment. **C)** Left, side view of the Dual-Stage Module filter featuring coconut fibers wrapped in two brims of pāpale lau niu inspired weaving, encasing a Baseline Prototype module inside for extra filtration. Middle, top view and cross section of the Dual-Stage Module. Right, example of water flow during deployment.

#### 2.2 Pāpale Lau Niu Brim construction

The pāpale lau niu brim used for the subsequent models was constructed using fresh coconut fronds (18 leaflets per side) which were cut, split lengthwise, and thinned by shaving down the central stalk until the leaflets were flexible enough for weaving. Each half was bent into a ring (∅ ≈ 25 cm) and secured with twine. Following the natural leaflet orientation, an over-one/under-one weave was carried around the circle twice, then tightened and trimmed, yielding a sturdy brim that dries to a rigid frame after one month of sun-curing.

#### 2.3 Single-Stage Module model construction

The “Single-Stage Module” model (Figure 3B) was constructed using the pāpale lau niu brim as follows. Two cured brims as woven in Section 2.2 served as “plates.” 10 g of dried coconut fiber were spread on one plate; the second plate was placed fiber-side-down, and their rims were laced together with jute. The remaining leaflets were tucked flat to create a dense, double-layer weave, resulting in a Single-Stage Module with a total mass of ≈ 150 g. Photographic steps of the module construction are provided in Supplementary Data 1.

#### 2.4 Dual-Stage Module model construction

The Dual-Stage Module shown in Figure 3C combines the buoyant husk sphere from the Baseline Prototype with twin coconut-leaf “crowns.” Each crown brim was woven as in Section 2.2, but four adjacent leaflets were pulled through the inner band to form a shallow bowl for surface flotation. After one-month of drying, the husk sphere was sandwiched between the crowns, and 20 g of coconut fiber were distributed in the annular space before the rims were tied. This dual-stage configuration adds vertical flow paths and increases buoyancy. Photographic steps of the module construction are provided in Supplementary Data 1.

#### 2.5 Testing sunscreen and plastic filtering efficiency

All filtering experiments were conducted in a tank with the following design. A 100 L clear storage tote was filled with 50 L of tap water containing 35 g/L NaCl at 25 °C. During each trial the container was gently rocked 5 cm back and forth at 1 Hz to simulate wave action.

To test for plastic removal, LDPE fragments of 2.5 mm each were made from Perler beads to represent microplastics. 300 LDPE fragments were dispersed and stirred in the container. Filter modules were tethered at the center of the tank using jute twine, allowing it to float at the air-water interface while preventing lateral drift and maintaining consistent positioning across the trials. The filters were subjected to the motion for 2 minutes. The module was retrieved, and retained particles were manually counted. The tank was strained with 1 mm x 1 mm mesh to remove residual plastics before the next run. Each filter module was tested in four consecutive trials using the same physical module, with the tank reset, residual microplastics removed, and fiber inside the modules replaced between trials. Statistical analyses for significant differences were performed by single-factor ANOVA.

To test for sunscreen removal, 100 mL of SPF 50 sunscreen (Coppertone) was whisked into a clean container set up as above, except string was used to divide the surface of the container into a 5x5 grid. The module was floated for 60 minutes under continuous rocking. At 0, 15, 30, 45, and 60 minutes, four 100 mL samples were collected from randomly generated grid coordinates, clarified with silicone-oiled turbidimeter cuvettes, and turbidity was recorded in triplicate for each sample. Identical sampling was then performed in sunscreen-free seawater with loose coconut fiber to quantify turbidity contributions from leached organics. Net sunscreen removal was calculated by comparing the turbidity of sunscreen-amended water surrounding the submerged filters with sunscreen-free control tubs, thereby accounting for turbidity changes caused by organic leachates released from the filter materials. Statistical analyses for significant differences were performed by single-factor ANOVA.

Three modules of the Dual-Stage Module design connected with jute twine were then deployed at a local beach on the southeastern shore of O‘ahu, Hawai‘i to test material integrity in a real ocean environment for one hour. Video of the deployed filters is available in Supplementary Data 1.

## Results

### Coconut husk naturally harbors fungi tolerant of and potentially degrading plastic- and sunscreen-associated compounds

To test whether the innate microbial community of coconut husks could grow on sunscreen and plastic compounds, we isolated fungi from the husks on plates with sunscreen or plastic as their main carbon source. Multiple fungal colonies grew faster on media amended with pollutants compared with non-amended controls (Figure 1, Supplementary Figure 1). Moreover, localized indentations formed in the sunscreen-containing medium surrounding some fungal colonies, indicating a physical change in the medium associated with fungal growth (Figure 1B), and some colonies formed a zone of clearance on plastic plates, indicating plastic degradation or modification (Figure 1D). In total we identified 24 isolates that could grow on either sunscreen or plastic media (Table 1). Sequencing of the internal transcribed region (ITS) identified 16 of those isolates to genus level, composing of genera *Chaetomium*, *Mrakia*, *Curvularia*, *Meyerozyma*, *Cladosporium*, and *Fusarium* (Table 1).

To confirm that these isolates could still grow with added pollutants in saline conditions, we chose four isolates to grow in plastic and sunscreen liquid media with added 35 g/L NaCl. All four fungal isolates showed growth both on control carbon-limited media with no added carbon source and on carbon-limited media supplemented with sunscreen or plastic. Three isolates appeared to favor growth on plastic over the control, however growth did not significantly differ between any treatments from any isolate (Figure 2).

### Three prototypes of filter devices woven with coconut materials differ in their ability to filter plastic and sunscreen

To utilize the filtering ability of coconut fibers, we tested three devices for easy deployment into aqueous environments (Figure 3). The first Baseline Prototype encases coconut fiber with dried coconut husks, held together with jute twine (Figure 3A). The following two designs—the Single-Stage Module and Dual-Stage Module—are based on the Hawaiian pāpale lau niu, an application of Polynesian weaving techniques that honors cultural heritage while shaping the system’s functional geometry (Figure 3B,C). By leveraging the natural properties of coconut materials, we expected the woven designs would enhance pollutant capture through several complementary mechanisms. Epicuticular wax on leaf surfaces could promote hydrophobic adhesion of sunscreen residues (Riedel et al., 2009), while tightly woven interstices may improve retention of suspended microplastics. In addition, the circular geometry of the leaf crowns may alter local water flow and generate gentle rotational currents, increasing contaminant capture in dynamic aquatic environments.

After filter design and construction, we measured microplastic and sunscreen filtration efficacies of each filter module by floating them in tanks of saltwater contaminated with controlled amounts of either microplastic pieces or sunscreen. Each filter demonstrated varying capacities for microplastic removal: in two minutes the Baseline Prototype caught an average of 11.08 ± 7.91% of the 300 submerged plastic pieces, the Single-Stage Module design caught 69.83 ± 13.84%, and the Dual-Stage Module design caught 76.58 ± 3.73% (Figure 4A,B). The Single-Stage Module (ANOVA, P=3.19×10^-4^) and Dual-Stage Module (ANOVA, P=5.56×10^-6^) captured significantly more particles than the Baseline Prototype. The Single-Stage and Dual-Stage module capture rates did not significantly differ.

**Figure 4.**
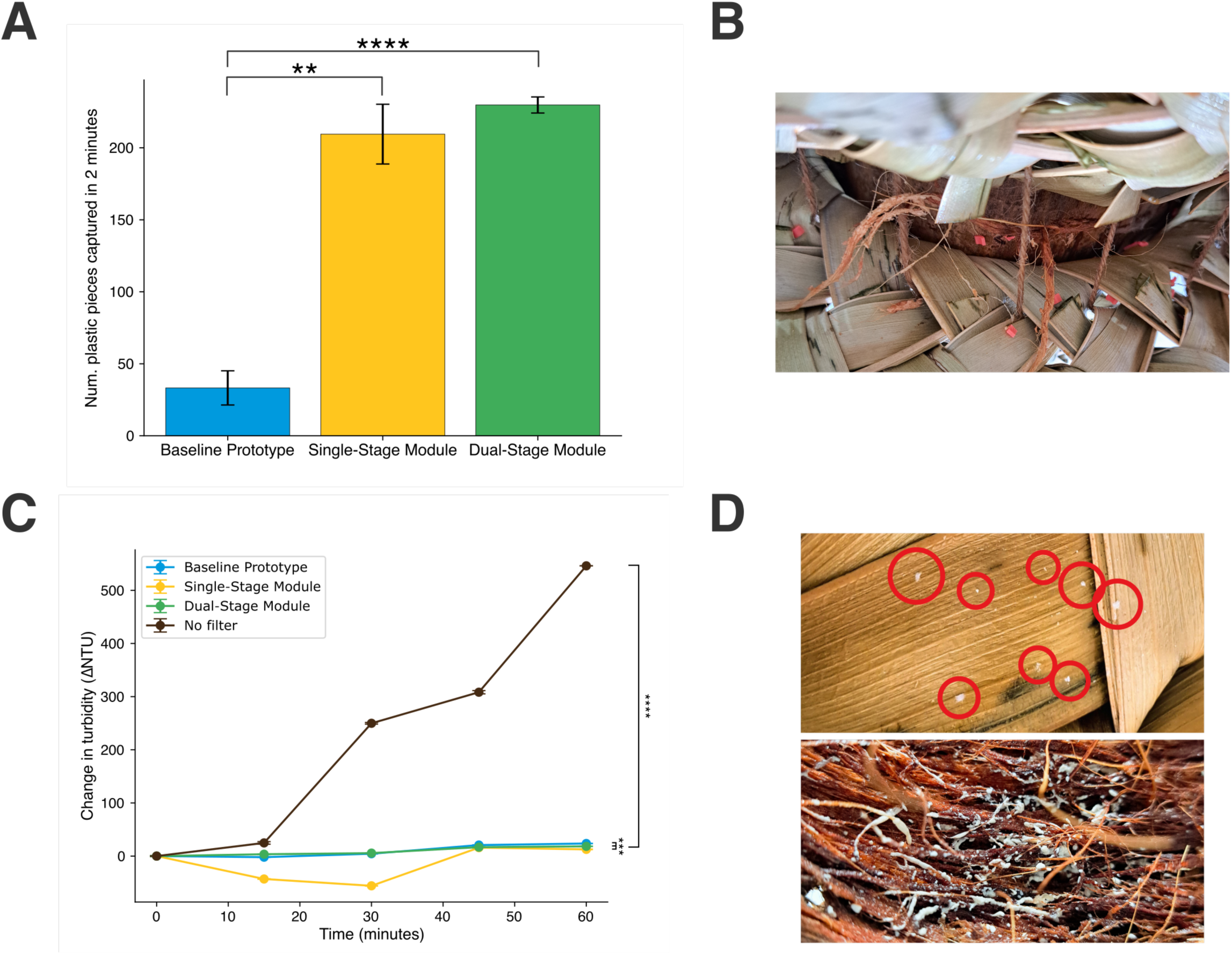
Each filter design is differentially effective in filtering plastic and sunscreen. **A)** Removal of microplastics after two minutes by woven filters. Error bars indicate standard error, with significant interactions denoted by asterisks. **B)** Photo of microplastics caught in the Dual-Stage filter after trials. **C)** Removal of sunscreen after 1 hour by woven filters. Error bars indicate standard error, with significant interactions denoted by asterisks. **D)** Photo of sunscreen particles that had adhered to the outer leaves of the Dual-Stage Module filter after the trials, circled in red, and of sunscreen trapped in the coconut fibers.

Each design also decreased the turbidity of surrounding sunscreen water compared to a no filter control. While the turbidity of the sunscreen inoculated water increased over time during each trial (likely due to increased clumping of the hydrophobic sunscreen), all three filter designs ended with lower turbidity measurements than the control trial with no filter. Each design showed varying ending turbidity, with the Baseline Prototype ending with a net change in turbidity 522.45 ± 0.41 NTU lower than the no filter control, the Single-Stage Module design ending with a 533.51 ± 0.81 NTU lower net turbidity, and the Dual-Stage Module design ending with a 527.79 ± 0.44 NTU lower net turbidity. All filter ending net turbidity changes were significantly lower than the no filter control (ANOVA, P=5.57×10^-19^). The Single-Stage Module significantly differed from the Baseline Module (ANOVA, P=2.97×10^-5^) and from the Dual-Stage Module (ANOVA, P=4.23×10^-4^). The Dual-Stage Module also significantly differed from the Baseline Module (ANOVA, P=1.05×10^-4^). Additionally, the Single-Stage Module design resulted in a unique large initial decrease in turbidity within 30 minutes, demonstrating a unique ability for fast initial capture of sunscreen (Figure 4C,D).

For in-field testing, three filters based on the Dual-Stage Module design were constructed and interconnected using jute twine, then deployed at a local beach east of Honolulu, Hawai‘i. Both microplastics and sunscreen-derived contaminants have previously been documented in coastal environments around O□ahu, particularly near the field testing site. For example: surveys at Makapu□u Beach detected approximately 199 microplastic particles m□² of beach sediment (Rey, Franklin & Rey, 2021), and sampling across five sites in Maunalua Bay detected oxybenzone at all locations, with one site reaching 19.2 µg/L (Downs et al., 2016). The filters maintained structural integrity and stability under natural coastal conditions throughout the one-hour testing period, and visible particles were trapped on the leaves of the Dual-Stage Module’s crown brim after removal (Figure 5).

**Figure 5.**
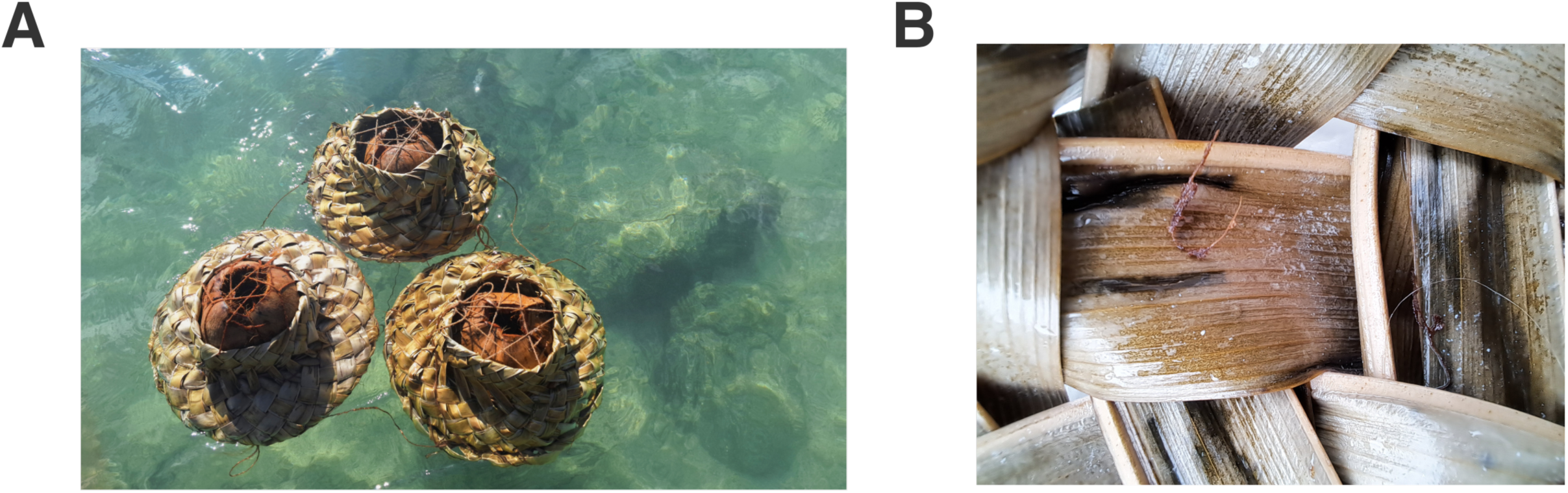
Three filters modeled after Dual-Stage Module deployed in an eastern Honolulu beach for field testing. **A)** Picture of the three filters, connected with jute twine, floating in the ocean for field testing. **B)** Picture of particles trapped in the filter device after removal from water.

## Discussion

With plastic and sunscreen pollution on the rise, an effective and easily implementable clean-up solution is of increasing need. This research expands on previous literature utilizing coconut fibers as biodegradable, cost-effective pollutant filters. We analyzed the potential role of naturally occurring microbes in pollution remediation, identifying 24 fungal isolates from coconut husks capable of growing under plastic- and/or sunscreen-containing conditions. For environmental deployment, we introduce a novel design to contain the coconut husks inspired by the traditional Hawaiian pāpale lau niu weaving.

While most remediation strategies focus primarily on physical pollutant removal, our results suggest that coconut-fiber filters may provide the additional benefit of supporting naturally occurring fungal communities with potential pollutant-transforming capabilities. We recovered multiple fungal isolates from coconut husks that were able to grow under sunscreen- and plastic-containing conditions, indicating that the filter material can naturally harbor microbes with traits relevant to bioremediation. The fungal isolates included *Chaetomium*, *Curvularia*, *Meyerozyma*, *Cladosporium*, and *Fusarium*, all genera containing members previously demonstrated to degrade synthetic plastic polymers (Vivi, Martins-Franchetti & Attili-Angelis, 2019; Ren et al., 2021; Lou et al., 2022; Bhanot et al., 2024; Sathiyabama et al., 2024). These reports span several polymer types, including polyethylene, polystyrene, polycaprolactone, and polyurethane, supporting the broader observation that coconut husks naturally harbor fungal taxa with established polymer-degrading potential. In contrast, direct degradation of sunscreen UV-filter compounds has not, to our knowledge, been demonstrated for these five genera, despite evidence that other fungi can transform organic UV-filter chemical compounds used in sunscreen such as oxybenzone (benzophenone-3) and 4-methylbenzylidene camphor (Badia-Fabregat et al., 2012). The growth of our coconut-associated isolates under sunscreen-containing conditions is therefore particularly noteworthy, although further research would be needed to establish degradation of the UV-filter compounds. Several of the identified genera possess broad metabolic or extracellular enzymatic capabilities relevant to the transformation of recalcitrant organic compounds; for example, *Chaetomium* is associated with lignocellulose decomposition (Longoni et al., 2012), while plastic degradation by *Cladosporium* and *Fusarium* has been linked to oxidative or hydrolytic enzymes including ligninolytic enzymes, esterases, and lipases (Ren et al., 2021; Sathiyabama et al., 2024). These characteristics provide a biologically plausible basis for further research of their ability to transform sunscreen-derived compounds. Additionally, our testing of a subset of the isolates in pollutant-containing liquid saline media supports the plausibility of these fungi surviving in oceanic conditions during deployment for use in bioremediation.

Each of our three filtration prototypes showed successful pollutant capture at different efficiencies. While the Single-Stage and Dual-Stage modules featuring the pāpale lau niu weaving outperformed the Baseline Prototype with just the coconut husks, the Dual-Stage module captured slightly more microplastics while the Single-Stage module absorbed sunscreen at a faster rate. Additionally, the Single-Stage module requires less materials and time to make, however the enhanced mass of the Dual-Stage module could better protect it from rough ocean conditions. More research is thus needed to deduce the best prototype for real-world application, however the survival of the Dual-Stage Module during our short-term field testing on O‘ahu shores provides promising support for further evaluation of the design through longer deployments, replicated trials, and direct chemical confirmation of pollutant removal.

With the combined findings of a natural sunscreen- and plastic-compatible microbial community and the high filtration efficiencies of our filter prototypes, we propose that the coconut husk functions as a dual-purpose platform for bioremediation: it acts as a physical filtration medium that captures sunscreen and plastic marine contaminants within its fibers, and also as a bioreactor that can enrich for fungi with potential abilities to degrade those same pollutants. Building on a recent study which demonstrates that artificially-created biofilms on coconut fibers enhance microplastic capture rates (Zharkenov et al., 2024), we thus envision a future filtration module which merges our designs with biofilms of pollutant-degrading microbes, either of naturally-occurring fungi as studied here or introduced, allowing trapped pollutant particles to be remineralized by the microbes in situ. In any case our woven filters, with or without a biofilm, could potentially be connected into larger modular chains for nearshore deployment in high-traffic beach areas, where they may help intercept sunscreen residues and suspended microplastics at local pollution hotspots.

The open burning of coconut husk biomass equivalent to that used in one Dual-Stage Module would release approximately 1.16 kg CO□, 0.93 g black carbon, and 5.4 g fine particulate matter (PM□.□) (Obeng et al., 2020). Repurposing this biomass in filtration modules could therefore reduce combustion-related emissions while simultaneously providing a biodegradable material for coastal pollutant remediation. This use of locally available plant waste for environmental remediation builds upon Indigenous Hawaiian perspectives emphasizing resource stewardship and reciprocal relationships with the environment (Kealiikanakaoleohaililani & Giardina, 2016). By integrating niu (coconut) materials, traditional weaving practices, physical pollutant capture, and naturally associated fungal communities, we hope that our work here provides a framework for further development of locally grounded plastic and sunscreen remediation strategies for Hawai‘i beaches and beyond.

## Acknowledgements

We gratefully acknowledge organization Livable Maunalua Hui for permitting the collection of coconuts and coconut fronds from the historic coconut tree grove of its managed wetlands for use in this study. We would also like to thank the Hach Water Equipment Grant program for providing instrumentation and equipment used for water-quality measurements of the sunscreen removal assays. Funding from the Hawai‘i Young Investigators Program supported DNA sequencing of fungal isolates, enabling taxonomic identification of the microbial communities examined in this study. We would also like to thank Renee Takara for laboratory training, technical guidance, and assistance with experimental techniques and the UH Manoa ASGPB Sequencing Lab for Sanger sequencing services. The first author is especially grateful to Ronja Steinbach for facilitating her introduction to the Amend Lab and connecting her with the mentorship and laboratory resources that enabled the microbiological component of this research.

## Supplementary figure captions

**Supplemental Figure 1.** Coconut husk extracted plated on control plates with no sunscreen or plastic added showing little to no fungal growth.

**Supplemental Figure 2.** Microbial communities isolated from visually distinct coconut-husk extract conditions exhibit different growth patterns on plastic- and sunscreen-containing media. **A)** Example of coconut husk producing a light-colored extract after soaking for 10 days. **B)** Fungal isolates obtained from light-extract coconut husk and subsequently grown on sunscreen-containing media. **C)** Fungal isolates obtained from light-extract coconut husk and subsequently grown on plastic-containing media. **D)** Example of coconut husk producing a dark-colored extract after soaking for 10 days. **E)** Fungal isolates obtained from dark-extract coconut husk and subsequently grown on sunscreen-containing media. **F)** Fungal isolates obtained from dark-extract coconut husk and subsequently grown on plastic-containing media.

## Supplementary data captions

**Supplemental data 1.** Video of making filter designs.

## References

Altschul SF, Gish W, Miller W, Myers EW, Lipman DJ. 1990. Basic local alignment search tool. Journal of Molecular Biology 215:403–410. DOI: 10.1016/S0022-2836(05)80360-2.

Badia-Fabregat M, Rodríguez-Rodríguez CE, Gago-Ferrero P, Olivares A, Piña B, Díaz-Cruz MS, Vicent T, Barceló D, Caminal G. 2012. Degradation of UV filters in sewage sludge and 4-MBC in liquid medium by the ligninolytic fungus *Trametes versicolor*. Journal of Environmental Management 104:114–120. DOI: 10.1016/j.jenvman.2012.03.039.

Berger T, Whitner S, Reher R, Amend AS. 2026. Multi-omics insights into the enzymatic degradation of polyurethane by marine fungi. Journal of Hazardous Materials: Plastics 2:100040. DOI: 10.1016/j.hazmp.2026.100040.

Bhanot V, Mamta, Gupta S, Panwar J. 2024. Phylloplane fungus *Curvularia dactyloctenicola* VJP08 effectively degrades commercially available PS product. Journal of Environmental Management 351:119920. DOI: 10.1016/j.jenvman.2023.119920.

Breakell T, Kowalski I, Foerster Y, Kramer R, Erdmann M, Berking C, Heppt MV. 2024. Ultraviolet Filters: Dissecting Current Facts and Myths. Journal of Clinical Medicine 13:2986. DOI: 10.3390/jcm13102986.

Bucci K, Tulio M, Rochman CM. 2020. What is known and unknown about the effects of plastic pollution: A meta-analysis and systematic review. Ecological Applications 30:e02044. DOI: 10.1002/eap.2044.

Cardoso CKM, Moreira ÍTA, Queiroz AF de S, Oliveira OMC de, Lobato AK de CL. 2025. Multiscale Evaluation of Raw Coconut Fiber as Biosorbent for Marine Oil Spill Remediation: From Laboratory to Field Applications. Resources 14:159. DOI: 10.3390/resources14100159.

Chong MY, Tam YJ. 2020. Bioremediation of dyes using coconut parts via adsorption: a review. SN Applied Sciences 2:187. DOI: 10.1007/s42452-020-1978-y.

Damikouka I, Anastasopoulou M, Vgenopoulou E. 2024. Sunscreens in the aquatic environment and potential solutions for mitigation of sunscreen pollution. Euro-Mediterranean Journal for Environmental Integration 9:1833–1850. DOI: 10.1007/s41207-024-00655-4.

Downs CA, Cruz OT, Remengesau Jr. TE. 2022. Sunscreen pollution and tourism governance: Science and innovation are necessary for biodiversity conservation and sustainable tourism. Aquatic Conservation: Marine and Freshwater Ecosystems 32:896– 906. DOI: 10.1002/aqc.3791.

Downs CA, Fauth JE, Halas JC, Dustan P, Bemiss J, Woodley CM. 2002. Oxidative stress and seasonal coral bleaching. Free Radical Biology and Medicine 33:533–543. DOI: 10.1016/S0891-5849(02)00907-3.

Downs CA, Kramarsky-Winter E, Segal R, Fauth J, Knutson S, Bronstein O, Ciner FR, Jeger R, Lichtenfeld Y, Woodley CM, Pennington P, Cadenas K, Kushmaro A, Loya Y. 2016. Toxicopathological Effects of the Sunscreen UV Filter, Oxybenzone (Benzophenone-3), on Coral Planulae and Cultured Primary Cells and Its Environmental Contamination in Hawaii and the U.S. Virgin Islands. Archives of Environmental Contamination and Toxicology 70:265–288. DOI: 10.1007/s00244-015-0227-7.

Ekanayaka AH, Tibpromma S, Dai D, Xu R, Suwannarach N, Stephenson SL, Dao C, Karunarathna SC. 2022. A Review of the Fungi That Degrade Plastic. Journal of Fungi 8:772. DOI: 10.3390/jof8080772.

Eriksen M, Cowger W, Erdle LM, Coffin S, Villarrubia-Gómez P, Moore CJ, Carpenter EJ, Day RH, Thiel M, Wilcox C. 2023. A growing plastic smog, now estimated to be over 170 trillion plastic particles afloat in the world’s oceans—Urgent solutions required. PLOS ONE 18:e0281596. DOI: 10.1371/journal.pone.0281596.

Falk-Andersson J, Rognerud I, De Frond H, Leone G, Karasik R, Diana Z, Dijkstra H, Ammendolia J, Eriksen M, Utz R, Walker TR, Fürst K. 2023. Cleaning Up without Messing Up: Maximizing the Benefits of Plastic Clean-Up Technologies through New Regulatory Approaches. Environmental Science & Technology 57:13304–13312. DOI: 10.1021/acs.est.3c01885.

Gregory MR. 2009. Environmental implications of plastic debris in marine settings— entanglement, ingestion, smothering, hangers-on, hitch-hiking and alien invasions. Philosophical Transactions of the Royal Society B: Biological Sciences 364:2013–2025. DOI: 10.1098/rstb.2008.0265.

Johnson B. 2024. Plastic-eating bacteria boost growing business of bioremediation. Nature Biotechnology 42:1481–1485. DOI: 10.1038/s41587-024-02401-1.

Kealiikanakaoleohaililani K, Giardina CP. 2016. Embracing the sacred: an indigenous framework for tomorrow’s sustainability science. Sustainability Science 11:57–67. DOI: 10.1007/s11625-015-0343-3.

Keawe LOMA, MacDowell M, Dewhurst CK. 2014. ‘Ike Ulana Lau Hala: The Vitality and Vibrancy of Lau Hala Weaving Traditions in Hawai‘i. University of Hawaii Press.

Lanctôt CM, Bednarz VN, Melvin S, Jacob H, Oberhaensli F, Swarzenski PW, Ferrier-Pagès C, Carroll AR, Metian M. 2020. Physiological stress response of the scleractinian coral Stylophora pistillata exposed to polyethylene microplastics. Environmental Pollution 263:114559. DOI: 10.1016/j.envpol.2020.114559.

Lee CE, Messer LF, Wattiez R, Matallana-Surget S. 2025. The invisible threats of sunscreen as a plastic co-pollutant: Impact of a common organic UV filter on biofilm formation and metabolic function in the nascent marine plastisphere. Journal of Hazardous Materials 495:139103. DOI: 10.1016/j.jhazmat.2025.139103.

Lima CLBS, Moreira ÍTA, Campos LMA, Pontes LAM, Teixeira LSG. 2025. Removal of arsenic from landfill leachate using green coconut fiber. Scientific Reports 15:37064. DOI: 10.1038/s41598-025-20861-6.

Longoni P, Rodolfi M, Pantaleoni L, Doria E, Concia L, Picco AM, Cella R. 2012. Functional Analysis of the Degradation of Cellulosic Substrates by a Chaetomium globosum Endophytic Isolate. Applied and Environmental Microbiology 78:3693–3705. DOI: 10.1128/AEM.00124-12.

Lou H, Fu R, Long T, Fan B, Guo C, Li L, Zhang J, Zhang G. 2022. Biodegradation of polyethylene by *Meyerozyma guilliermondii* and *Serratia marcescens* isolated from the gut of waxworms (larvae of *Plodia interpunctella*). Science of The Total Environment 853:158604. DOI: 10.1016/j.scitotenv.2022.158604.

Lozano C, Givens J, Stien D, Matallana-Surget S, Lebaron P. 2020. Bioaccumulation and Toxicological Effects of UV-Filters on Marine Species. In: Tovar-Sánchez A, Sánchez-Quiles D, Blasco J eds. Sunscreens in Coastal Ecosystems: Occurrence, Behavior, Effect and Risk. Cham: Springer International Publishing, 85–130. DOI: 10.1007/698_2019_442.

Mason VG, Skov MW, Hiddink JG, Walton M. 2022. Microplastics alter multiple biological processes of marine benthic fauna. Science of The Total Environment 845:157362. DOI: 10.1016/j.scitotenv.2022.157362.

Mendrik FM, Henry TB, Burdett H, Hackney CR, Waller C, Parsons DR, Hennige SJ. 2021. Species-specific impact of microplastics on coral physiology. Environmental Pollution 269:116238. DOI: 10.1016/j.envpol.2020.116238.

Nikiema J, Asiedu Z. 2022. A review of the cost and effectiveness of solutions to address plastic pollution. Environmental Science and Pollution Research 29:24547–24573. DOI: 10.1007/s11356-021-18038-5.

Nurtayeva A, Rakhmonov J, Sarykova A, Rachana K, Kristanti RA. 2025. Microbial Biodegradation of Sunscreen Agents: Mechanisms, Enzymatic Pathways, and Environmental Implications. Tropical Aquatic and Soil Pollution 5:167–184. DOI: 10.53623/tasp.v5i2.824.

Obeng GY, Amoah DY, Opoku R, Sekyere CKK, Adjei EA, Mensah E. 2020. Coconut Wastes as Bioresource for Sustainable Energy: Quantifying Wastes, Calorific Values and Emissions in Ghana. Energies 13:2178. DOI: 10.3390/en13092178.

Pinheiro HT, MacDonald C, Santos RG, Ali R, Bobat A, Cresswell BJ, Francini-Filho R, Freitas R, Galbraith GF, Musembi P, Phelps TA, Quimbayo JP, Quiros TEAL, Shepherd B, Stefanoudis PV, Talma S, Teixeira JB, Woodall LC, Rocha LA. 2023. Plastic pollution on the world’s coral reefs. Nature 619:311–316. DOI: 10.1038/s41586-023-06113-5.

Pogosa J. 2018. Productivity and Sustainability of Coconut Production and Husk Utilization in the Philippines: Coconut Husk Availability and Utilization.

Ren GC, Pang AM, Gao Y, Wu SX, Ge ZQ, Zhang TF, Wanasinghe DN, Khan S, Mortimer PE, Xu JC, Gui H, Ren GC, Pang AM, Gao Y, Wu SX, Ge ZQ, Zhang TF, Wanasinghe DN, Khan S, Mortimer PE, Xu JC, Gui H. 2021. Polyurethane-degrading fungi from soils contaminated with rocket propellant and their ability to decompose alkyne terminated polybutadiene with urethane. Studies in Fungi 6:224–239. DOI: 10.5943/sif/6/1/15.

Rey SF, Franklin J, Rey SJ. 2021. Microplastic pollution on island beaches, Oahu, Hawaìi. PLoS ONE 16:e0247224. DOI: 10.1371/journal.pone.0247224.

Riedel M, Riederer M, Becker D, Herran A, Kullaya A, Arana-López G, Peña-Rodríguez L, Billotte N, Sniady V, Rohde W, Ritter E. 2009. Cuticular wax composition in Cocos nucifera L.: physicochemical analysis of wax components and mapping of their QTLs onto the coconut molecular linkage map. Tree Genetics & Genomes 5:53–69. DOI: 10.1007/s11295-008-0168-7.

Rodríguez-Romero A, Ruiz-Gutiérrez G, Viguri JR, Tovar-Sánchez A. 2019. Sunscreens as a New Source of Metals and Nutrients to Coastal Waters. Environmental Science & Technology 53:10177–10187. DOI: 10.1021/acs.est.9b02739.

Sánchez-Quiles D, Blasco J, Tovar-Sánchez A. 2020. Sunscreen Components Are a New Environmental Concern in Coastal Waters: An Overview. In: Tovar-Sánchez A, Sánchez-Quiles D, Blasco J eds. Sunscreens in Coastal Ecosystems: Occurrence, Behavior, Effect and Risk. Cham: Springer International Publishing, 1–14. DOI: 10.1007/698_2019_439.

Sathiyabama M, Boomija RV, Sathiyamoorthy T, Mathivanan N, Balaji R. 2024. Mycodegradation of low-density polyethylene by Cladosporium sphaerospermum, isolated from platisphere. Scientific Reports 14:8351. DOI: 10.1038/s41598-024-59032-4.

Shaath N (ed.). 2005. Sunscreens: Regulations and Commercial Development. Boca Raton: CRC Press. DOI: 10.1201/b14170.

Steinbach RM, Whitner S, Amend AS. 2025. Marine fungi degrade plastic and can be conditioned to do it faster. Mycologia 117:1–8. DOI: 10.1080/00275514.2024.2422598.

Stelte W, Reddy N, Barsberg S, Sanadi A. 2022. Coir from coconut processing waste as a raw material for applications beyond traditional uses. BioResources 18. DOI: 10.15376/biores.18.1.Stelte.

Thushari GGN, Senevirathna JDM. 2020. Plastic pollution in the marine environment. Heliyon 6. DOI: 10.1016/j.heliyon.2020.e04709.

Vivi VK, Martins-Franchetti SM, Attili-Angelis D. 2019. Biodegradation of PCL and PVC: Chaetomium globosum (ATCC 16021) activity. Folia Microbiologica 64:1–7. DOI: 10.1007/s12223-018-0621-4.

Wang T, Li B, Shi H, Ding Y, Chen H, Yuan F, Liu R, Zou X. 2024. The processes and transport fluxes of land-based macroplastics and microplastics entering the ocean via rivers. Journal of Hazardous Materials 466:133623. DOI: 10.1016/j.jhazmat.2024.133623.

White T, Bruns T, Lee S, Taylor J, Innis M, Gelfand D, Sninsky J. 1990. Amplification and Direct Sequencing of Fungal Ribosomal RNA Genes for Phylogenetics. In: Pcr Protocols: a Guide to Methods and Applications,. 315–322.

Zharkenov Y, Mkilima T, Abduova A, Zhaksylykova L, Turashev A, Imambayeva R, Imambaev N, Jaxymbetova M, Smagulova A, Beysenbaeva E. 2024. Utilizing biofilm-enhanced coconut coir for microplastic removal in wastewater. Case Studies in Chemical and Environmental Engineering 9:100726. DOI: 10.1016/j.cscee.2024.100726.

