## Supplementary figures and images for "Coconut filters woven using Native Hawaiian techniques capture coastal microplastic and sunscreen pollution and support fungal bioremediation"

### Supplementary figures

#### Supplementary figure 1

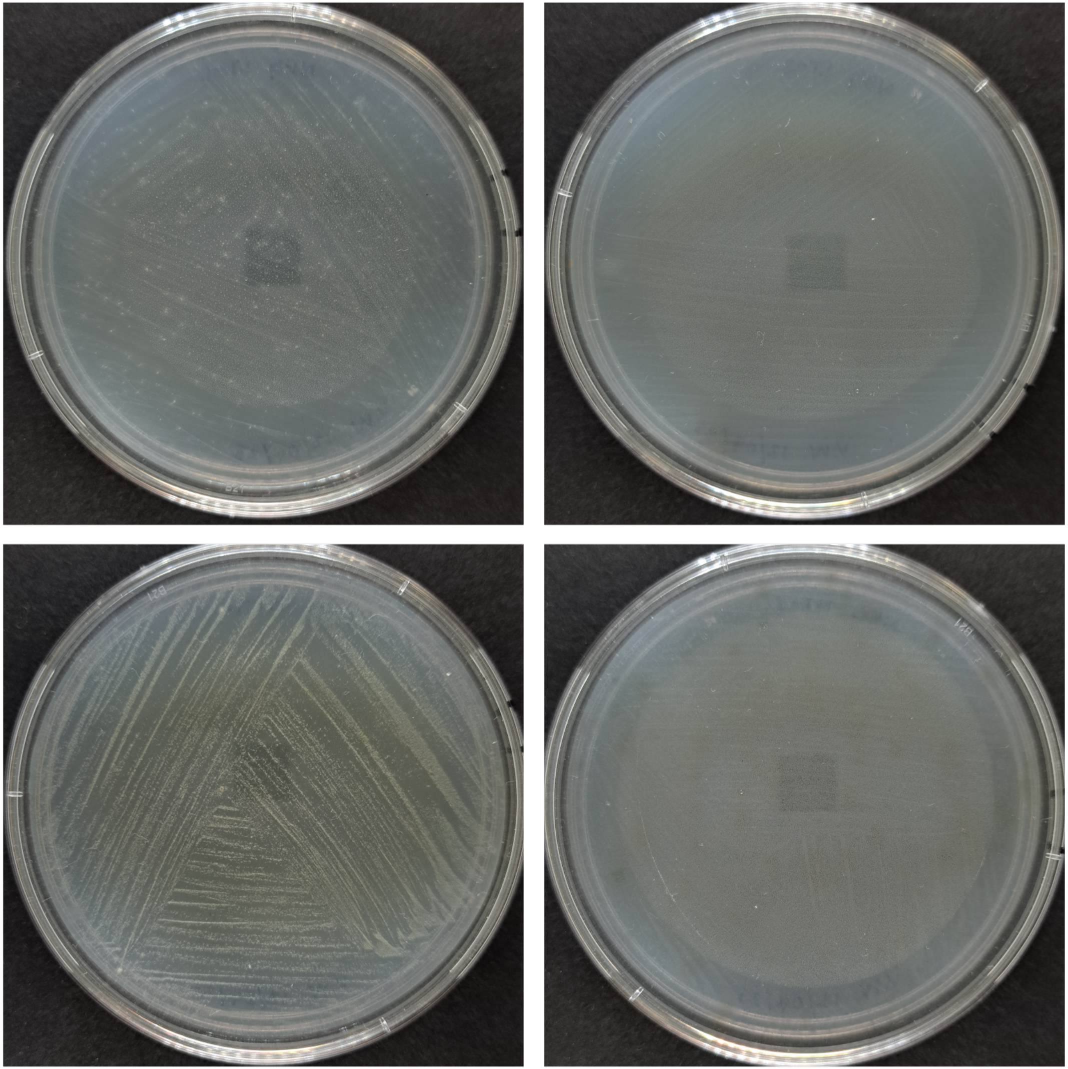

#### Supplementary figure 2

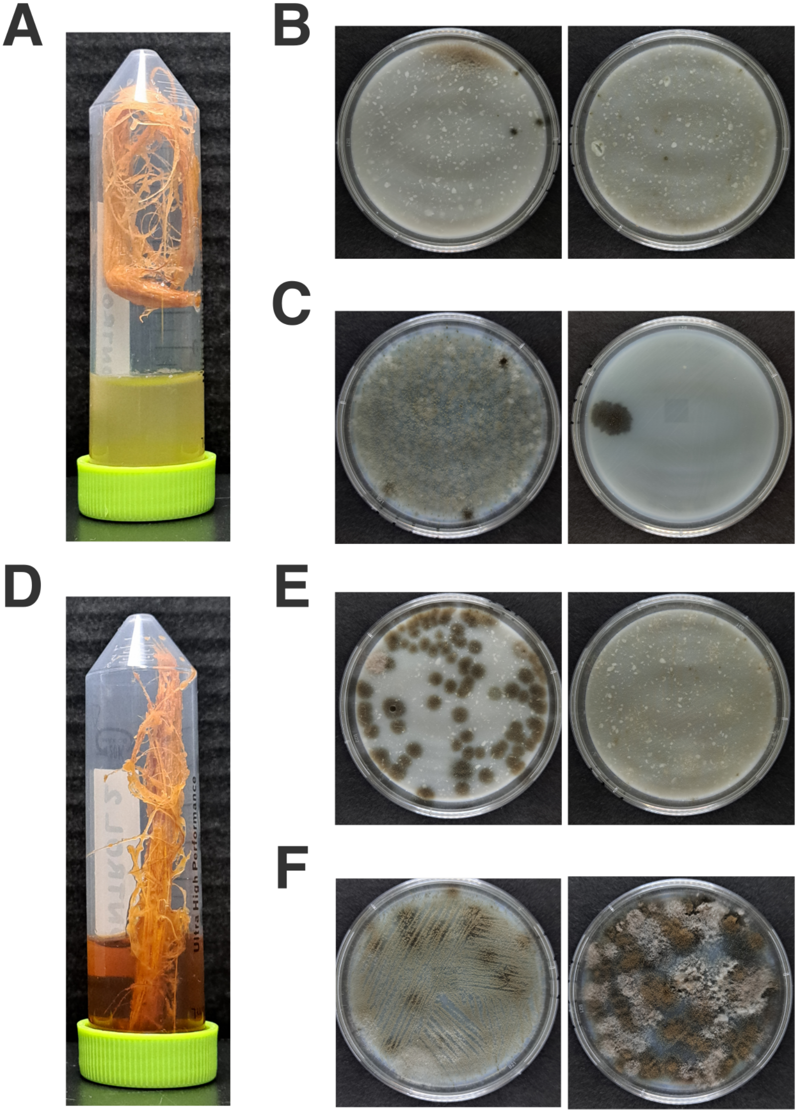
